# A SARS CoV-2 nucleocapsid vaccine protects against distal viral dissemination

**DOI:** 10.1101/2021.04.26.440920

**Authors:** Jacob Class, Tanushree Dangi, Justin M. Richner, Pablo Penaloza-MacMaster

**Author notes:** **Correspondence:** Justin Richner, Pablo Penaloza-MacMaster.

## Abstract

The SARS CoV-2 pandemic has killed millions of people. This viral infection can also result in substantial morbidity, including respiratory insufficiency and neurological manifestations, such as loss of smell and psychiatric diseases. Most SARS CoV-2 vaccines are based on the spike antigen, and although they have shown extraordinary efficacy at preventing severe lung disease and death, they do not always confer sterilizing immune protection. We performed studies in K18-hACE2 mice to evaluate whether the efficacy of SARS CoV-2 vaccines could be augmented by incorporating nucleocapsid as a vaccine antigen. We vaccinated mice with adenovirus-based vaccines encoding spike antigen alone, nucleocapsid antigen alone, or combined spike and nucleocapsid antigens. Mice were then challenged intranasally with SARS CoV-2, and acute viral loads were quantified at a proximal site of infection (lung) and a distal site of infection (brain). Interestingly, the spike-based vaccine conferred acute protection in the lung, but not in the brain. The spike-based vaccine conferred acute protection in the brain only if combined with the nucleocapsid-based vaccine. These findings suggest that nucleocapsid-specific immunity is important for the distal control of SARS CoV-2, warranting the inclusion of nucleocapsid in next-generation COVID-19 vaccines.

## Introduction

Most SARS CoV-2 vaccines used in humans are based on the spike antigen. These vaccines have shown high efficacy against severe disease and death, but they do not always confer sterilizing immunity^1–6^. In particular, breakthrough infections can be detected in nasopharyngeal swabs of vaccinated individuals^7–9^. Nasopharyngeal swabs typically contain viruses derived from the proximal site of challenge (the respiratory system), which may not reflect ongoing virus replication at distal sites of the body. It is unknown whether SARS CoV-2 vaccines can prevent acute viral dissemination to distal sites of the body, such as the central nervous system, which is considered an immune-privileged site that is relatively “impermeable” to circulating antibodies^10^.

The central nervous system has become increasingly important in the management of COVID-19 disease, especially because even mild or asymptomatic infections can trigger neurological manifestations, including “brain fog” and psychiatric conditions^11^. SARS CoV-2 is a respiratory virus, but prior reports suggest that it may also disseminate to the brain via the olfactory mucosa^12–14^. Post-mortem analyses of COVID-19 patients have shown the presence of SARS CoV-2 in the brain^12,15–17^. Knowing whether vaccines can block viral dissemination to the central nervous system is important, because this would help elucidate if vaccinated people who get exposed to the virus could still develop neurological complications. Due to ethical reasons it is not feasible to sample the brain of vaccinated individuals to assess breakthrough infections in this distal site.

The SARS CoV-2 spike protein is considered to be a critical antigen target in coronavirus vaccines. This antigen alone is the basis for the Pfizer/BioNTek vaccine, Moderna vaccine, Johnson & Johnson’s (J&J) Janssen vaccine, AstraZeneca vaccine, CanSino vaccine, Sputnik V vaccine, Novavax vaccine, among others. Vaccines that encode the viral spike generate robust neutralizing antibodies that prevent the initial entry of SARS CoV-2 into the respiratory system, but it is still unclear whether other viral antigens provide equally important immune protection. It is also unclear whether inclusion of multiple viral antigens could provide a synergistic improvement in vaccine-elicited protection. In this study, we compared the efficacy of spike-based versus nucleocapsid-based vaccines after an intranasal SARS CoV-2 challenge in K18-hACE2 mice, which are highly susceptible to SARS CoV-2 infection. Surprisingly, we show that a spike-based vaccine does not provide acute protection to the central nervous system; protection against distal viral dissemination to the nervous system was only observed when a spike-based vaccine was co-administered with a nucleocapsid-based vaccine. These findings demonstrate a potent synergy between spike-specific and nucleocapsid-specific immune responses, and provide a framework for the rational design of coronavirus vaccines.

## Results

### Immunogenicity of spike-based and nucleocapsid-based SARS CoV-2 vaccines

K18-hACE2 mice constitute a robust pre-clinical model that recapitulates salient features of SARS CoV-2 infection in humans, including respiratory insufficiency, a dysregulated inflammatory response, neurological complications and death. Therefore, this mouse model has been useful for investigating antiviral therapies, vaccines, and for understanding COVID-19 pathogenesis^18–22^. We first vaccinated K18-hACE2 mice intramuscularly with an adenovirus vector expressing: SARS CoV-2 spike (Ad5-S), nucleocapsid (Ad5-N), or both (Ad5-S+N), at a dose of 10^9^ PFU per mouse.

After three weeks, mice were bled. Peripheral blood mononuclear cells (PBMCs) were used for tetramer binding assays (K^b^VL8 to detect spike-specific CD8 T cells) or intracellular cytokine staining assays (ICS) to detect nucleocapsid-specific CD8 T cells; and sera were used for antibody quantification by enzyme-linked immunosorbent assay (ELISA) (Figure 1A). Ad5-S and Ad5-N vaccination elicited detectable CD8 T cell and antibody responses (Figure 1B-1E). These results confirm the immunogenicity of the spike-based and nucleocapsid-based vaccines.

**Figure 1.**
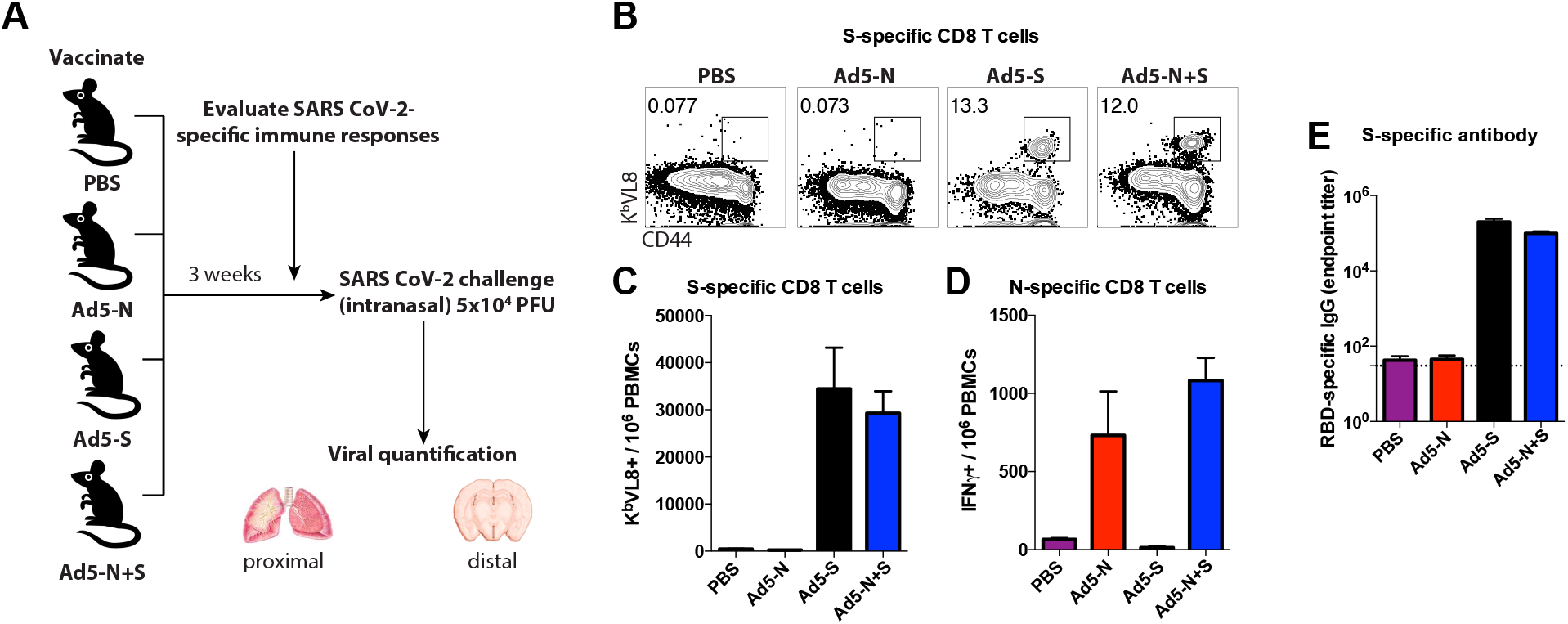
Immunogenicity of spike and nucleocapsid-based vaccines. **(A)** Experimental approach for evaluating immune responses and immune protection in K18-hACE2 mice. **(B)** Representative FACS plots showing the frequencies of SARS CoV-2-specific CD8 T cells (K^b^ VL8+) in PBMCs. **(C)** Summary of SARS CoV-2 spikespecific CD8 T cells (K^b^ VL8+) in PBMCs. **(D)** Summary of SARS CoV-2 nucleocapsid-specific CD8 T cells in PBMCs. To detect nucleocapsid-specific CD8 T cells we stimulated splenocytes for 5 hr with overlapping peptide pools (ICS). **(E)** Summary of SARS CoV-2 receptor binding domain (RBD)-specific antibody responses in sera. Data are from week 3 post-vaccination. Data are from an experiment with n=5 per group. Error bars represent SEM.

### A spike-based vaccine provides proximal protection in the lung, but not distal protection in the brain

We performed intranasal challenges with 5×10^4^ PFU of SARS CoV-2 (isolate USA-WA1/2020), followed by euthanasia 72 hr post-challenge to evaluate acute viral loads in tissues. We also challenged unvaccinated K18-hACE2 mice as controls. First, we compared viral loads in the lung, which represents a proximal site of challenge. Intranasal challenge of unvaccinated K18-hACE2 mice with SARS CoV-2 normally results in high viral replication in lung, followed by distal dissemination to the brain by day 2 post-challenge^18^. Consistent with prior studies^18–21^, the unvaccinated mice showed high viral loads in lung (Figure 2A). Mice vaccinated with the nucleocapsid-based vaccine showed a pattern of improved antiviral control in lung, but this was not statistically significant. In contrast, mice vaccinated with the spike-based vaccine exhibited significant antiviral protection in the lung (Figure 2A).

**Figure 2.**
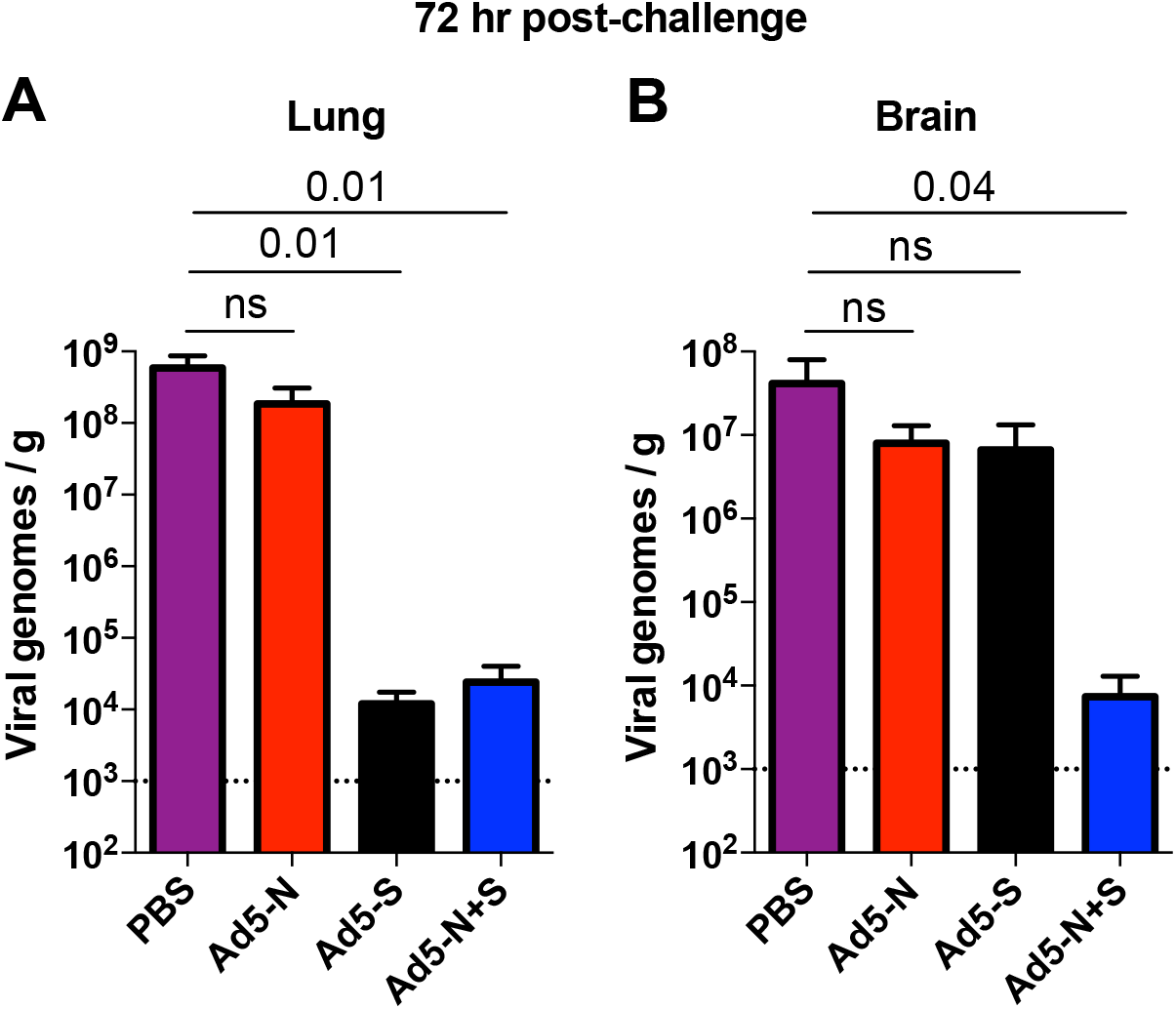
Immune protection of spike and nucleocapsid-based vaccines. RNA was harvested from the lungs **(A)** and brains **(B)** at 72 hr post-infection, and viral RNA was quantified by RT-qPCR. Data are from an experiment with n=5 per group. Error bars represent SEM. Indicated P-values were calculated using Kruskal-Wallis test (Dunn’s multiple comparisons). NS, not significant (p>0.05).

Importantly, vaccination of mice with both spike-based and nucleocapsid-based vaccines did not confer any synergistic protective advantage in the lung, relative to spike-based vaccine alone (Figure 2A). These data show that a spike-based vaccine alone is sufficient to protect the site of initial viral entry, the respiratory system.

### Effect of nucleocapsid-specific immunity in the distal control of a SARS CoV-2 infection

At first glance, the data above suggested that nucleocapsid-specific immunity plays a dispensable role during a SARS CoV-2 infection. Nucleocapsid-specific immunity has recently garnered attention as a main target for T cell responses^23–25^. Antibody responses block the initial entry of viruses at proximal sites of challenge, but once the infection has occurred, T cell responses are critical for controlling second-round infections and subsequent viral dissemination to distal sites. SARS CoV-2 is thought to disseminate to the nervous system after second-round infections that progress from the olfactory mucosa to the brain^14^. Although current SARS CoV-2 vaccines show high efficacy in protecting the respiratory system via their induction of neutralizing antibodies, it is not clear whether they can provide acute protection at distal sites.

We show that mice immunized with a spike-based vaccine (or a nucleocapsid-based vaccine) exhibit a pattern or reduced viral loads in brain relative to unimmunized mice, but this was not statistically significant (Figure 2B). However, combining a spike-based vaccine with a nucleocapsid-based vaccine resulted in significantly improved protection in brain (Figure 2B). Taken together, these findings show that spike-specific immunity confers acute protection in the site of initial viral entry (the respiratory system), but acute protection in distal sites may critically depend on also having nucleocapsid-specific immunity.

## Discussion

Several SARS CoV-2 vaccines have received emergency use authorization (EUA), including the Pfizer/BioNTek vaccine, Moderna vaccine, Johnson & Johnson’s (J&J) Janssen vaccine, AstraZeneca/Oxford vaccine, CanSino vaccine, Sputnik V vaccine, and Novavax vaccine. All of these vaccines are based solely on the spike protein of SARS CoV-2, and have demonstrated high efficacy against severe COVID-19. Although these vaccines prevent severe disease and death, breakthrough infections can occur in vaccinated individuals, suggesting that vaccine efficacy can be further improved.

Insidious neurological manifestations are common after SARS CoV-2 infection, and it is unknown if vaccines can block acute SARS CoV-2 replication in the nervous system. An unanticipated result from our studies is that a spike-based vaccine does not protect the central nervous system early following infection. This observation could help explain symptomatic infections that are reported in a fraction of vaccinated individuals, which can be associated with loss of smell and other neurological manifestations.

The establishment of initial infection foci in the respiratory system is inhibited by spike-specific antibodies, which are secreted at the mucosa and can block binding of the virus to the ACE2 receptor. On the other hand, secondary infection foci in distal tissues may be more susceptible to cytotoxic T cells, due to their ability to transmigrate into tissues and kill virally-infected cells. The nucleocapsid protein of SARS CoV-2 has been suggested to be an important target for T cell responses^23–25^. Firstly, this protein contains conserved cross-reactive T cell epitopes that are present among different coronaviruses, suggesting that it could be a target for universal coronavirus vaccines^26^. Secondly, the nucleocapsid protein is among the most abundant structural proteins in the coronavirus lifecycle^27–29^, which may facilitate antigen presentation and recognition by T cells.

A limitation of our study is that we only measured viral loads at a very early timepoint (72 hr post-challenge). We did not compare viral loads at other time points, because unvaccinated animals succumb by day 6 post-challenge^19^, and because it is known that spike-based vaccination elicits immune responses that ultimately clear SARS CoV-2 within a week of challenge^30,31^. Our study was focused on evaluating acute viral control at a very early time post-challenge. Future studies will determine whether these findings generalize to other vaccine platforms besides adenovirus-based vectors (e.g. mRNA vaccines encoding nucleocapsid), and whether other virus-specific immune responses (e.g. envelope-, membrane-specific) can improve further distal protection. Although the K18-hACE2 model is considered useful to evaluate vaccines and antivirals, the extensive acute viral dissemination observed in brain may be a peculiarity of this mouse model. Nevertheless, it is still reasonable to conclude that nucleocapsid-specific immunity improves distal viral control of SARS CoV-2. In conclusion, we show a substantial benefit of including both spike and nucleocapsid antigens in COVID-19 vaccines. These results warrant the incorporation of nucleocapsid as a critical antigen in future coronavirus vaccines.

## Materials and Methods

### Mice and vaccinations

6-8-week-old K18-hACE2 mice were used. These mice express the human ACE2 protein behind the keratin 18 promoter, directing expression in epithelial cells. Mice were purchased from Jackson laboratories (Stock No: 034860). Approximately half were males and half were females. Mice were immunized intramuscularly (50 μL per quadriceps) with an Ad5 vector expressing SARS CoV-2 spike protein (Ad5-S), or nucleocapsid protein (Ad5-N), or both; diluted in sterile PBS, at 10^9^ PFU per mouse. Ad5-N was a kind gift of the Masopust/Vezys laboratory^32^. These are non-replicating Ad5 vectors (E1/E3 deleted). The vectors contain a CMV (Cytomegalovirus) promoter driving the expression of the respective proteins. The Ad5 vectors were propagated on trans-complementing HEK293 cells (ATCC), purified by cesium chloride density gradient centrifugation, titrated, and then frozen at −80 °C.

### SARS CoV-2 virus and infections

The following reagent was deposited by the Centers for Disease Control and Prevention and obtained through BEI Resources, NIAID, NIH: SARS-Related Coronavirus 2, Isolate USA-WA1/2020, NR-52281. Virus was propagated and tittered on Vero-E6 cells. K18-hACE2 mice were anesthetized with isoflurane and challenged with 5×10^4^ PFU of virus. Mouse infectious were performed at the University of Illinois at Chicago (UIC) following BL3 guidelines with approval by the UIC Institutional Animal Care and Use Committee (IACUC).

### SARS CoV-2 quantification in tissues

Tissues were isolated from infected mice and homogenized in sterile PBS. RNA was isolated with the Zymo 96-well RNA isolation kit (Catalog #: R1052) following the manufacturer’s protocol. SARS CoV-2 viral burden was measured by qRT-PCR using Taqman primer and probe sets from IDT with the following sequences: Forward 5’ GAC CCC AAA ATC AGC GAA AT 3’, Reverse 5’ TCT GGT TAC TGC CAG TTG AAT CTG 3’, Probe 5’ ACC CCG CAT TAC GTT TGG TGG ACC 3’. A SARS CoV-2 copy number control was obtained from BEI (NR-52358) and used to quantify SARS-CoV-2 genomes.

### Reagents, flow cytometry and equipment

Single cell suspensions were obtained from PBMCs. Dead cells were gated out using Live/Dead fixable dead cell stain (Invitrogen). SARS CoV-2 nucleocapsid peptide pools were obtained through BEI Resources, NIAID, NIH (Peptide Array, SARS-Related Coronavirus 2 Nucleocapsid (N) Protein, NR-52404). These peptide pools were used for intracellular cytokine staining (ICS) to detect nucleocapsid-specific CD8 T cells. The peptide stimulation was performed in the presence of GolgiPlug and GolgiStop (BD Biosciences) for 5 hr at 37°C in a 5% CO_2_ incubator. MHC class I monomers (K^b^VL8, VNFNFNGL) were used for detecting spike-specific CD8 T cells, and were obtained from the NIH tetramer facility located at Emory University. MHC monomers were tetramerized in-house. The VNFNFNGL epitope is located in position 539-546 of the SARS CoV-2 spike protein, or position 525-532 of the SARS CoV-1 spike protein. Cells were stained with fluorescently-labeled antibodies against CD8α (53-6.7 on PerCP-Cy5.5), CD44 (IM7 on Pacific Blue), K^b^VL8 or IFNγ (XMG1.2) on APC. Fluorescently-labeled antibodies were purchased from BD Pharmingen, except for anti-CD44 (which was from Biolegend). Flow cytometry samples were acquired with a Becton Dickinson Canto II or an LSRII and analyzed using FlowJo (Treestar).

### SARS CoV-2-specific ELISA

Binding antibody titers were quantified using ELISA as described previously ^33,34^, using spike protein as a coating antigen. In brief, 96-well flat bottom plates MaxiSorp (Thermo Scientific) were coated with 0.1μg/well of SARS CoV-2 RBD protein, for 48 hr at 4°C. Plates were washed with PBS +0.05% Tween-20. Blocking was performed for 4 hr at room temperature with 200 μL of PBS + 0.05% Tween-20 + bovine serum albumin. 6μL of sera were added to 144 μL of blocking solution in first column of plate, 1:3 serial dilutions were performed until row 12 for each sample and plates were incubated for 60 minutes at room temperature. Plates were washed three times followed by addition of goat anti-mouse horseradish peroxidase-conjugated IgG (Southern Biotech) diluted in blocking solution (1:5000), at 100 μL/well and incubated for 60 minutes at room temperature. Plates were washed three times and 100 μL /well of Sure Blue substrate (Sera Care) was added for approximately 8 minutes. Reaction was stopped using 100 μL/well of KPL TMB stop solution (Sera Care). Absorbance was measured at 450 nm using a Spectramax Plus 384 (Molecular Devices). SARS CoV-2 RBD protein was made at the Northwestern Recombinant Protein Production Core by Dr. Sergii Pshenychnyi using a plasmid that was produced under HHSN272201400008C and obtained through BEI Resources, NIAID, NIH: Vector pCAGGS Containing the SARS-Related Coronavirus 2, Wuhan-Hu-1 Spike Glycoprotein Gene (soluble, stabilized), NR-52394.

### Statistical analysis

Statistical analyses are indicated on the figure legend. Dashed lines in data figures represent limit of detection. Statistical significance was established at p ≤0.05. Data were analyzed using Prism (Graphpad).

## Findings

Spike-based vaccination protects the lung, but does not prevent acute viral dissemination to the brain.

A spike-based vaccine together with a nucleocapsid-based vaccine confers acute protection in lung and brain.

## References

1 Polack, F. P. et al. Safety and Efficacy of the BNT162b2 mRNA Covid-19 Vaccine. N Engl J Med 383, 2603–2615, doi:10.1056/NEJMoa2034577 (2020).

2 Baden, L. R. et al. Efficacy and Safety of the mRNA-1273 SARS-CoV-2 Vaccine. N Engl J Med 384, 403–416, doi:10.1056/NEJMoa2035389 (2021).

3 Sadoff, J. et al. Safety and Efficacy of Single-Dose Ad26.COV2.S Vaccine against Covid-19. The New England journal of medicine, doi:10.1056/NEJMoa2101544 (2021).

4 Voysey, M. et al. Safety and efficacy of the ChAdOx1 nCoV-19 vaccine (AZD1222) against SARS-CoV-2: an interim analysis of four randomised controlled trials in Brazil, South Africa, and the UK. Lancet 397, 99–111, doi:10.1016/S0140-6736(20)32661-1 (2021).

5 Zhu, F. C. et al. Immunogenicity and safety of a recombinant adenovirus type-5-vectored COVID-19 vaccine in healthy adults aged 18 years or older: a randomised, double-blind, placebo-controlled, phase 2 trial. Lancet 396, 479–488, doi:10.1016/s0140-6736(20)31605-6 (2020).

6 Logunov, D. Y. et al. Safety and efficacy of an rAd26 and rAd5 vector-based heterologous prime-boost COVID-19 vaccine: an interim analysis of a randomised controlled phase 3 trial in Russia. Lancet 397, 671–681, doi:10.1016/s0140-6736(21)00234-8 (2021).

7 Tande, A. J. et al. Impact of the COVID-19 Vaccine on Asymptomatic Infection Among Patients Undergoing Pre-Procedural COVID-19 Molecular Screening. Clinical infectious diseases: an official publication of the Infectious Diseases Society of America, doi:10.1093/cid/ciab229 (2021).

8 Keehner, J. et al. SARS-CoV-2 Infection after Vaccination in Health Care Workers in California. The New England journal of medicine, doi:10.1056/NEJMc2101927 (2021).

9 Levine-Tiefenbrun, M. et al. Initial report of decreased SARS-CoV-2 viral load after inoculation with the BNT162b2 vaccine. Nature medicine, doi:10.1038/s41591-021-01316-7 (2021).

10 Forrester, J. V., McMenamin, P. G. & Dando, S. J. CNS infection and immune privilege. Nature reviews. Neuroscience 19, 655–671, doi:10.1038/s41583-018-0070-8 (2018).

11 Nalbandian, A. et al. Post-acute COVID-19 syndrome. Nat Med 27, 601–615, doi:10.1038/s41591-021-01283-z (2021).

12 Song, E. et al. Neuroinvasion of SARS-CoV-2 in human and mouse brain. J Exp Med 218,doi:10.1084/jem.20202135 (2021).

13 Cantuti-Castelvetri, L. et al. Neuropilin-1 facilitates SARS-CoV-2 cell entry and infectivity. Science 370, 856–860, doi:10.1126/science.abd2985 (2020).

14 Meinhardt, J. et al. Olfactory transmucosal SARS-CoV-2 invasion as a port of central nervous system entry in individuals with COVID-19. Nat Neurosci 24, 168–175, doi:10.1038/s41593-020-00758-5 (2021).

15 Kumari, P. et al. Neuroinvasion and Encephalitis Following Intranasal Inoculation of SARS-CoV-2 in K18-hACE2 Mice. Viruses 13,doi:10.3390/v13010132 (2021).

16 Meinhardt, J. et al. Olfactory transmucosal SARS-CoV-2 invasion as a port of central nervous system entry in individuals with COVID-19. Nature neuroscience 24, 168–175, doi:10.1038/s41593-020-00758-5 (2021).

17 Gasmi, A. et al. Neurological Involvements of SARS-CoV2 Infection. Molecular neurobiology 58, 944–949, doi:10.1007/s12035-020-02070-6 (2021).

18 Winkler, E. S. et al. SARS-CoV-2 infection of human ACE2-transgenic mice causes severe lung inflammation and impaired function. Nat Immunol 21, 1327–1335, doi:10.1038/s41590-020-0778-2 (2020).

19 Oladunni, F. S. et al. Lethality of SARS-CoV-2 infection in K18 human angiotensin-converting enzyme 2 transgenic mice. Nat Commun 11,6122, doi:10.1038/s41467-020-19891-7 (2020).

20 Rosenfeld, R. et al. Post-exposure protection of SARS-CoV-2 lethal infected K18-hACE2 transgenic mice by neutralizing human monoclonal antibody. Nat Commun 12,944, doi:10.1038/s41467-021-21239-8 (2021).

21 Zheng, J. et al. COVID-19 treatments and pathogenesis including anosmia in K18-hACE2 mice. Nature 589, 603–607, doi:10.1038/s41586-020-2943-z (2021).

22 Bao, L. et al. The pathogenicity of SARS-CoV-2 in hACE2 transgenic mice. Nature 583, 830–833, doi:10.1038/s41586-020-2312-y (2020).

23 Rydyznski Moderbacher, C. et al. Antigen-Specific Adaptive Immunity to SARS-CoV-2 in Acute COVID-19 and Associations with Age and Disease Severity. Cell 183, 996–1012 e1019, doi:10.1016/j.cell.2020.09.038 (2020).

24 Nguyen, T. H. O. et al. CD8+ T cells specific for an immunodominant SARS-CoV-2 nucleocapsid epitope display high naïve precursor frequency and T cell receptor promiscuity. Immunity, doi:https://doi.org/10.1016/j.immuni.2021.04.009 (2021).

25 Lineburg, K. E. et al. CD8+ T cells specific for an immunodominant SARS-CoV-2 nucleocapsid epitope cross-react with selective seasonal coronaviruses. Immunity, doi:https://doi.org/10.1016/j.immuni.2021.04.006 (2021).

26 Dutta, N. K., Mazumdar, K. & Gordy, J. T. The Nucleocapsid Protein of SARS-CoV-2: a Target for Vaccine Development. J Virol 94,doi:10.H28/JVI.00647-20 (2020).

27 de Breyne, S. et al. Translational control of coronaviruses. Nucleic Acids Res 48,1250212522, doi:10.1093/nar/gkaa1116 (2020).

28 Chang, C. K., Hou, M. H., Chang, C. F., Hsiao, C. D. & Huang, T. H. The SARS coronavirus nucleocapsid protein--forms and functions. Antiviral Res 103, 39–50, doi:10.1016/j.antiviral.2013.12.009 (2014).

29 Weiss, S. R. & Leibowitz, J. L. Coronavirus pathogenesis. Adv Virus Res 81, 85–164, doi:10.1016/B978-0-12-385885-6.00009-2 (2011).

30 Feng, L. et al. An adenovirus-vectored COVID-19 vaccine confers protection from SARS-COV-2 challenge in rhesus macaques. Nat Commun 11, 4207, doi:10.1038/s41467-020-18077-5 (2020).

31 Wu, S. et al. A single dose of an adenovirus-vectored vaccine provides protection against SARS-CoV-2 challenge. Nat Commun 11, 4081, doi:10.1038/s41467-020-17972-1 (2020).

32 Joag, V. et al. Cutting Edge: Mouse SARS-CoV-2 Epitope Reveals Infection and Vaccine-Elicited CD8 T Cell Responses. J Immunol 206, 931–935, doi:10.4049/jimmunol.2001400 (2021).

33 Dangi, T., Chung, Y. R., Palacio, N. & Penaloza-MacMaster, P. Interrogating Adaptive Immunity Using LCMV. CurrProtoc Immunol 130,e99, doi:10.1002/cpim.99 (2020).

34 Palacio, N. et al. Early type I IFN blockade improves the efficacy of viral vaccines. J Exp Med 217, doi:10.1084/jem.20191220 (2020).

